# Local urothelial cell-driven detrusor contractions

**DOI:** 10.1101/2025.01.28.635145

**Authors:** Firoj Alom, Kyle Joseph Ratcliff, Aaron D Mickle

## Abstract

Signaling molecules released from the urothelium by mechanical stretch are known to play a sensory role in bladder contractions via neuronal signaling. It is also theorized that these molecules released from the urothelium could act locally to induce urothelial cell-mediated local bladder contractions. In this study, we specifically stimulated the urothelial cells using optogenetics to investigate how the urothelial released signaling molecules can influence bladder contractions locally. Using an *ex-vivo* whole bladder preparation, we stimulated the urothelial cells by activating channelrhodopsin-2 (ChR2) with blue light and initiated the urothelial cell-mediated local bladder contractions. The P2X receptor antagonist, PPADS, nearly abolished the contractions. The muscarinic receptor antagonist, atropine, significantly inhibited the contractions. Nifedipine, which blocks extracellular Ca^2+^ entry, abolished the contractions. G protein-coupled receptor inhibitor YM-254890 significantly inhibited the contractions. Hemichannel inhibitor carbenoxolone disodium and the exocytotic pathway of transmitter release inhibitor brefeldin A also showed significant inhibition of the contractions. In conclusion, we validated the previous hypothesis that urothelial release factors can influence bladder contractions locally without the need for signaling from the central nervous system. Further studies are needed to determine the relevance of this signaling pathway in normal bladder physiology and pathophysiologic conditions.

## Introduction

The innermost layer of the bladder, the urothelium, contributes to sensory signaling and communicates environmental changes to other associated tissues, including nerve endings and the detrusor muscle^1,2^. Recently, using a transgenic mouse model, we showed that direct stimulation of the urothelial cells results in pelvic nerve afferent firing, which leads to bladder contractions^3^. Additionally, in *ex vivo* bladder preparations, we showed that stimulation of the urothelial cells can also produce a smaller bladder contraction without an intact CNS^3^. These findings led us to conduct the present study to explore the signaling mechanisms involved in the urothelial cell-mediated local bladder contractions.

Spontaneous contractile activity was observed in the urothelium-intact muscle preparations and disappeared when the mucosa (including the urothelium) was removed, suggesting that the urothelium influenced this spontaneous detrusor activity^4^. The intraluminal application of ATP induced contractions in isolated intact bladder preparations were significantly inhibited by atropine, and the removal of the urothelium caused a similar reduction of contractions as elicited by atropine^5^. Together, these results suggest that the urothelium can influence detrusor contractions.

Understanding the precise role of the urothelium in facilitating local bladder contractions remains a complex challenge. This is partly due to the presence of ATP and acetylcholine (Ach) receptors, the primary mediators of the detrusor contractions in the urothelium, afferent nerve endings in the lamina propria, and the detrusor muscle. Isolating the specific contribution of the urothelium in local contractions using a pharmacological approach is complicated by this distribution of receptors. Furthermore, the close proximity of these cell types and their capacity to release similar signaling molecules adds to the difficulty of this work.

To overcome these limitations, we developed a mouse model where ChR2, a light-activated non-selective cation channel, is selectively expressed in the urothelial cells. This urothelial-specific ChR2 mouse model allowed us to specifically stimulate the urothelial cells with light and study its downstream effects on the detrusor muscle. Using an *ex vivo* bladder preparation, we showed that the urothelial cells can evoke local bladder contractions by releasing ATP and ACh that act on the detrusor muscles. The signaling mechanism for these urothelial cell-mediated local bladder contractions seems to activate the existing pathways of purinergic and muscarinic receptors, leading to the contractions of the detrusor smooth muscles. Our approach in this study will have a long-lasting impact on biomedical research regarding how we study the urothelial-to-detrusor muscle communication and the changes that may occur under disease conditions.

## Materials and Methods

### Animals

All experiments were approved by the University of Florida Institutional Animal Care and Use Committee (IACUC). Experiments and mice handling were done in strict accordance with the National Institutes of Health (NIH) *Guide for the Care and Use of Laboratory Animals*. Male mice aged 9-17 weeks old were used in this study, and all experiments were performed during the light cycle (0600–1800 hours). Mice were housed on a 12:12-hour light-dark cycle and provided food and water ad libitum. Transgenic mice were produced by crossing the Ai32 line (No. 024109, The Jackson Laboratory) that expresses ChR2 in the presence of Cre, with uroplakin II (UPK2)-cre mice (No. 029281, The Jackson Laboratory) that expresses cre recombinase in the bladder urothelium^3,6,7^. This transgenic mouse line expresses ChR2 within the urothelium and allows for the targeted activation of the urothelial cells with exposure to blue light. These mice will be referred to as UPK2-ChR2 throughout the manuscript. We showed in our recently published paper that no overt differences were observed in optogenetic-evoked bladder contractions between male and female mice^3^. We used male mice in this study due to the technical advantage of securing fiber optics through the urethra compared to female mice, as males have a longer remaining urethra adjacent to the bladder neck.

### Tissue Preparation

Following exposure of mice to 2-3% isoflurane and confirmation of anesthetic plane, mice were quickly euthanized with cervical dislocation. The urinary bladders were quickly excised and placed in a petri dish filled with Kreb’s solution containing (in mM) 119 NaCl, 4.7 KCl, 24 NaHCO_3_, 1.2 KH_2_PO_4_, 2.5 CaCl_2_, 1.2 MgSO_4_, and 11 glucose at pH 7.3-7.4. The urinary bladder was freed from the connective and adipose tissues. The optical fiber (FT200UMT, Thor Labs) was inserted through the opening of the remaining urethra and secured with a 5-0 nylon suture (No. 07-809-8813, Patterson Veterinary). The prepared whole bladder was taken for isometric tension recording. The two ends of the bladder (dome and neck) were tied with 4-0 silk suture (07-891-0230, Patterson Veterinary) where a loop of thread was made at the bottom end (dome) to allow the tissue to be mounted to the isometric recording system **(Figure 1a)**.

**Figure 1.**
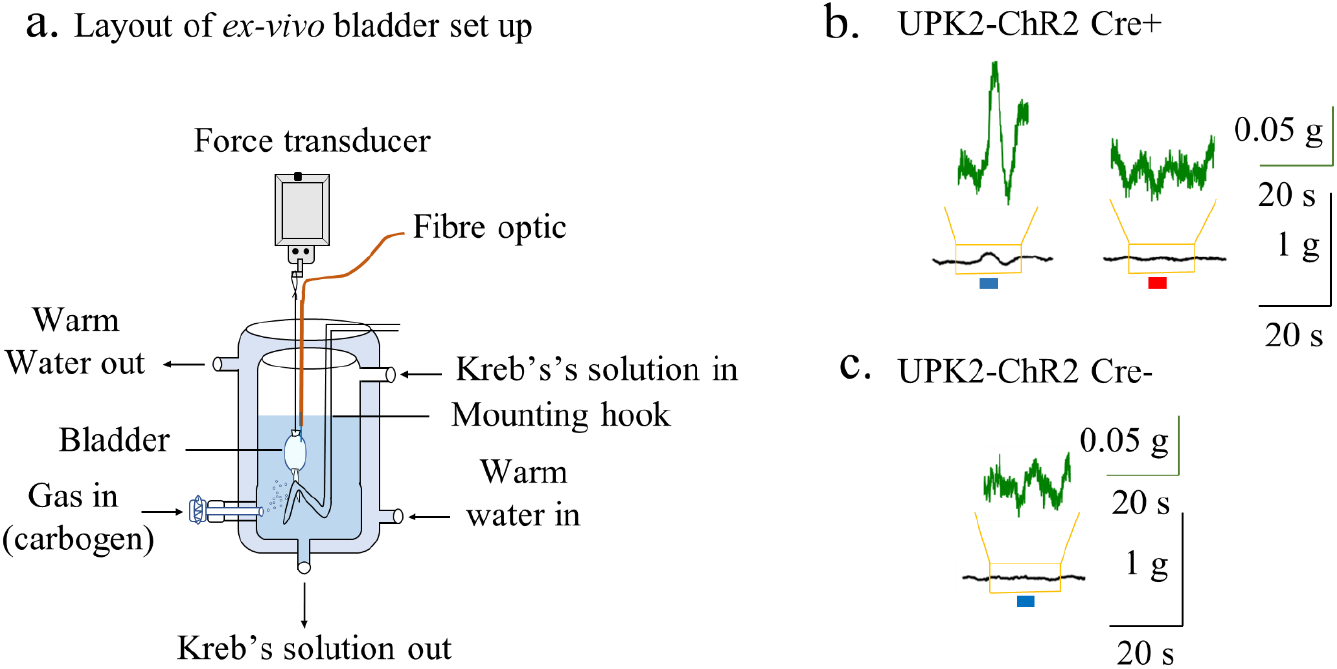
Ex-vivo bladder setup and recording the urothelial cell-mediated local bladder contractions. (a) An outline of the *ex vivo* whole bladder isometric tension recording setup. The prepared bladder was set up in a 10-mL organ bath containing Kreb’s solution bubbled with carbogen, and temperature (37°C) was maintained by using a circulating hot water bath. A fiber optic was inserted into the bladder through the remaining urethra and secured with nylon thread. One end of the bladder was attached to a glass hook, and the other to an isometric force transducer with silk thread to record the contractions in response to light or drugs. (b) Representative recording of the urothelial cell-mediated local bladder contractions evoked by blue light (20 mW, 5 s) and red light (20 mW, 5 s), showed that only the blue light activates the ChR2 expressed into the urothelial cells in UPK2-ChR2 Cre+ mice and produces contractions. (c) UPK2-ChR2 Cre-mice that do not express ChR2 into the urothelial cells did not produce contractions in response to blue light (20 mW, 5 s). Blue or red bars correspond to light stimulation.

### Isometric Tension Recording

The isometric tension recording of the bladder was performed as a modified version of the previously described^8^. The optical fiber was connected to a 473 nm blue laser (BL473T8-150FC, Lasercentury). The prepared bladder was vertically mounted in a 10-mL organ bath (Radnoti organ bath system, RSBRAD504/C) filled with Kreb’s solution and continuously aerated with 95% O_2_ and 5% CO_2_. The loop made at the dome of the bladder was hooked with a glass rod, and the other end was attached to a force transducer (FORT10G, World Precision Instruments) (**Figure 1a**). The transducer was attached with a Transbridge amplifier (TBM4, World Precision Instruments). The output of the amplifier was captured at a sampling rate of 1 kHz using a Micro 1401 data acquisition system (Cambridge Electronic Design) and Spike2 software (V10, Cambridge Electronic Design).

The bladders were subjected to ∼0.4-0.5 g of tension and allowed to equilibrate for 1 hour in fresh oxygenated Kreb’s solution. The bladder chamber temperature was maintained at 37°C with a circulating hot water bath (T72244004, Radnoti thermal bath/circulator).

After a one-hour equilibration period, we confirmed that bladders were viable and produced stable contractions by briefly (20-30 s) exposing the bladders to the hypertonic 70 mM KCl 3–5 times at 10–15 minutes intervals. After getting 2-3 consistent contractions with 70 mM KCl we applied 20 mW of blue light (5 s duration) 3 times at 2–3-minute intervals and induced the urothelial cell-mediated local bladder contractions. After establishing baseline bladder contractions, we directly applied different antagonists into the bath. After an incubation period (15–30 minutes), light was applied again 3 times to evoke the urothelial cell-mediated local bladder contractions. These light-evoked contractile responses were averaged and normalized relative to the corresponding high KCl contractions in the same bladder preparation.

### Drugs and Dilutions

Pyridoxalphosphate-6-azophenyl-2’,4’-disulfonic acid tetrasodium salt (PPADS) (0625) and adenosine-5’- (γ-thio)-triphosphate tetralithium salt (ATP) (4080) were purchased from Tocris Bioscience. Carbachol (10217179) and atropine sulphate (AAA1023609) were obtained from ThermoFisher Scientific. Nifedipine (4013408), carbenoxolone disodium (102624936), brefeldin A (0000216670), and YM-254890 (0000244228) were purchased from Sigma Aldrich. PPADS, ATP, carbachol, atropine sulphate and carbenoxolone disodium were diluted with saline water, whereas nifedipine, brefeldin A, and YM-254890 were diluted with dimethyl sulfoxide (DMSO) (4101628926). The final concentrations of the DMSO for these drugs varies from 0.0001% to 0.01%. The diluted drugs were aliquoted and stored at 4°C or -20°C according to manufacturer instructions. The following final concentrations of drugs used were PPADS (2 mM), atropine (2 µM), YM-254890 (100 nM), nifedipine (2 µM), carbenoxolone disodium (100 µM) and brefeldin A (200 µM). 3.5 M KCl stock solution was prepared and stored in 4°C to record high KCl (70 mM)-induced contractions.

### Data Analysis

The peak contractions produced in response to light, muscarinic agonist carbachol, and purinergic agonist ATP were expressed relative to the percentage of the reproducible 70 mM KCl-induced contractions measured in the same bladder preparation. The inhibitory percentage of drugs for the urothelial cell-mediated local bladder contractions was determined relative to the percentage of the control contractions before the addition of drugs into the bath solutions. Data are expressed as the mean ± standard error of the mean (SEM), n = the number of animals used. The Student’s paired t-test (two-tailed) was performed to determine the statistical significance of differences. The Shapiro-Wilk normality (Gaussian) test was performed to determine whether the distribution of values is parametric or non-parametric by using GraphPad Prism version 10.0.2. If the values passed the normality test, then a parametric Student’s paired t-test was conducted. Otherwise, a non-parametric Student’s paired t-test was conducted. Differences were considered statistically significant when p < 0.05. The original traces and column graphs were constructed by using GraphPad Prism software (version 10.1.1), and the figures were designed using Affinity Designer version 2.2.1.

## RESULTS

### Optogenetic Activation of the Urothelial Cells Elicits Local Bladder Contractions *ex vivo*

We have previously demonstrated that the application of blue light (20 mW, 5 s) produces a urothelial cell-mediated local bladder contraction in excised bladders as measured by intraluminal pressure^3^. In the present study, we showed the urothelial cell-mediated local bladder contractions specifically in *ex vivo* using an isometric force transducer from the excised bladder of UPK2-ChR2 mice. The urothelial cell-mediated local bladder contractions produced by light can clearly be distinguished from the baseline spontaneous contractions (**Figure 1b**). We also applied red light in the same Cre+ UPK2-ChR2 mice (**Figure 1b)** and blue light in Cre-UPK2-ChR2 mice (**Figure 1c**), which produced no contractions. These results suggest that these contractions produced by activation of the urothelial cells are specific to blue light that depends on the expression of ChR2 into the urothelial cells. The urothelial cell-mediated local bladder contractions relative to the 70 mM KCl-evoked contractions in the same preparations were 2.804 ± 0.28% (n = 35).

There were no significant differences (p = 0.128, n = 10) between the initial 70 mM KCl-evoked contractions and the final 70 mM KCl-evoked contractions after completion of light-evoked contractile studies with drugs (PPADS, atropine, YM-254890, carbenoxolone and brefeldin A). However, the 70 mM KCl-evoked contractions before and after the experiments with nifedipine were largely reduced.

### Effects of Non-specific P2 Purinergic Receptor Inhibitor PPADS on the Urothelial Cell-mediated Local Bladder Contractions

In this study, we investigated the role of ATP in producing urothelial cell-mediated local bladder contractions. Andersson, 2015 reviewed the role of ATP in activating purinergic receptors in inducing contractions of the bladder^9^. In our experiments, the urothelial cell-mediated local bladder contractions were significantly and almost completely inhibited by purinergic receptor inhibitor PPADS (2 mM) (**Figure 2a, 2b**). The urothelial cell-mediated local bladder peak contractions relative to 70 mM KCl-evoked contractions before and after application of PPADS (2 mM) were 1.91 ± 0.32% and 0.09 ± 0.08% (n = 4), respectively (**Figure 2b**). PPADS (2 mM) was able to inhibit optically activated contractile responses by 96.00 ± 3.01% (n = 4) when calculated as relative to percentage of control peak contractile responses (**Figure 2c**). These results confirmed that purinergic receptors are critical in inducing the urothelial cell-mediated local bladder contractions, as the contractions were mostly ablated by their inhibition.

**Figure 2.**
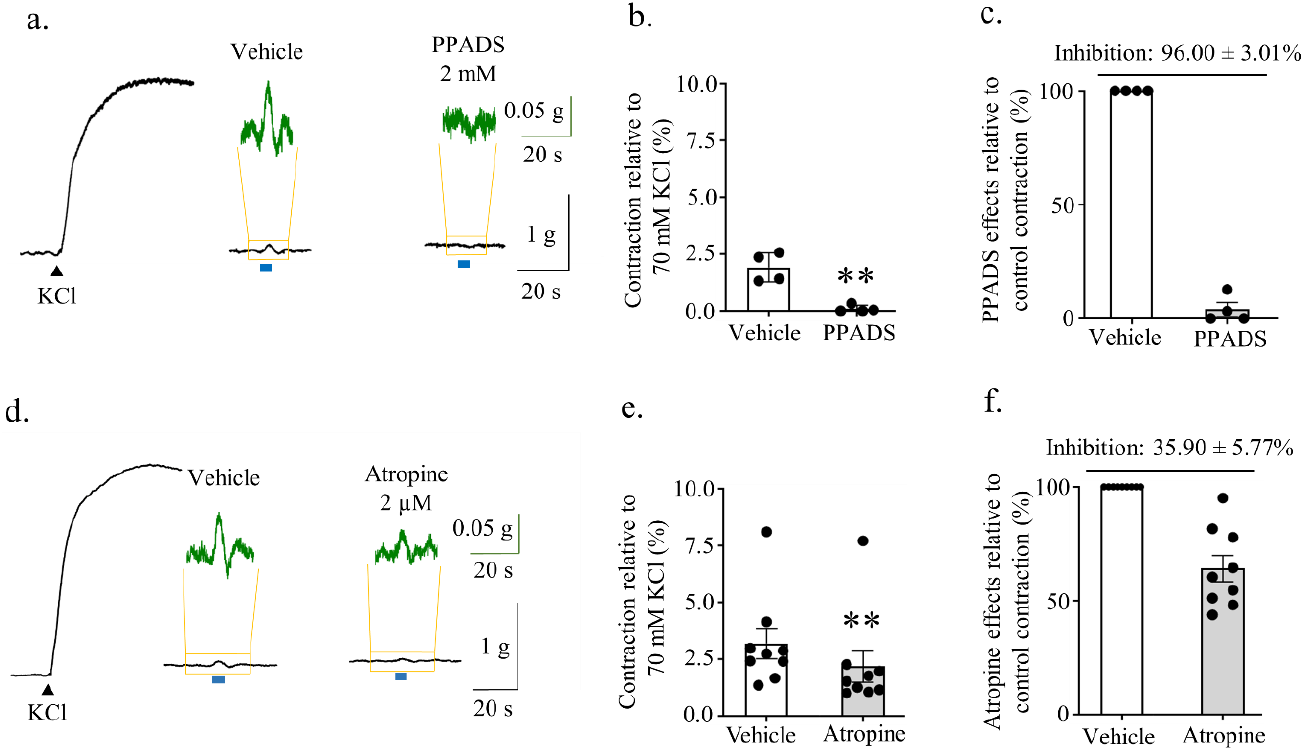
Effects of purinergic receptor inhibitor PPADS and muscarinic receptor antagonist atropine on the urothelial cell-mediated local bladder contractions. (a, d) Contractions evoked by 70 mM KCl and blue light (20 mW, 5 s). Blue light evoked urothelial cell-mediated local bladder contractions were recorded in the presence of vehicle and PPADS (2 mM) and atropine (2 µM), respectively. (b, e) Summarized data showing the inhibitory effects of PPADS (2 mM) and atropine (2 µM) on the urothelial cell-mediated local bladder contractions, respectively. (c, f) Summarized data of the percentage of inhibition of the urothelial cell-mediated local bladder contractions by PPADS (2 mM) and atropine (2 µM), respectively. The amplitude of the urothelial cell-mediated local bladder contractions was expressed as a percentage of the 70 mM KCl-evoked contractions in the same preparation. Each point represents the mean ± SEM (n = 4 for PPADS and n = 9 for atropine). **Significantly different (p = 0.007) from the corresponding control values. Blue bars correspond to light stimulation.

### Effects of Muscarinic Receptor Inhibitor Atropine on the Urothelial Cell-mediated Local Bladder Contractions

Previous findings showed that contractions evoked by the application of exogenous ATP inside the bladder lumen were inhibited by muscarinic receptor inhibitor atropine in *ex vivo* whole bladder preparation of rat^5^. These authors proposed that ATP may have a role in mediating the release of acetylcholine from the urothelial cells. Therefore, we investigated the possibility of whether the urothelial cell-mediated local bladder contractions involve muscarinic signaling pathways. Atropine (2 µM), a muscarinic receptor antagonist, significantly inhibited the urothelial cell-mediated local bladder contractions (**Figure 2d, 2e**). The mean peak amplitudes of the urothelial cell-mediated local bladder contractions relative to the 70 mM KCl-evoked contractile amplitudes before and after the application of atropine (2 µM) were 3.19 ± 0.67% and 2.19 ± 0.70% (n = 9), respectively (**Figure 2e**). The percentage of inhibition of contractions in the presence of atropine (2 µM) was 35.90 ± 5.77% (n = 9) relative to the corresponding control contractions (**Figure 2f**). These results suggest that muscarinic receptors also contribute to inducing the urothelial cell-mediated local bladder contractions.

### Effects of G-protein Coupled Receptor Inhibitor YM-254890 on the Urothelial-cell Mediated Local Bladder Contractions

G-protein coupled receptor (GPCR) signaling is activated through various receptors (M3, P2Y, H1, PGE2, etc.), which are involved in the contractions of the detrusor smooth muscles^10–13^. Here we investigated whether GPCR signaling is important for producing the urothelial cell-mediated local bladder contractions. YM-254890, is a broad-spectrum inhibitor for Gq and Gs proteins and shows a biased inhibition on G_i/o_ signaling^14^. YM-254890 (1 µM) was shown to abolish histamine (200 µM), and carbachol (200 nM) evoked contractions in the mouse bladder, indicating the involvement of histamine and muscarinic receptors coupling to Gq_/11_^12^. In our study, YM-254890 (100 nM) significantly inhibited the urothelial cell-mediated local bladder contractions (**Figure 3a, 3b**). The amplitudes of the peak contractions before and after the application of YM-254890 (100 nM) were 3.37 ± 0.74% and 2.09 ± 0.40% (n = 6) (**Figure 3b**), respectively. The inhibitory percentage of the contractions by YM-254890 (100 nM) was 33.89 ± 7.42% (n = 6) relative to control (vehicle) values (**Figure 3c**). These results suggest that GPCR signaling pathways are involved in producing the urothelial cell-mediated local bladder contractions.

**Figure 3.**
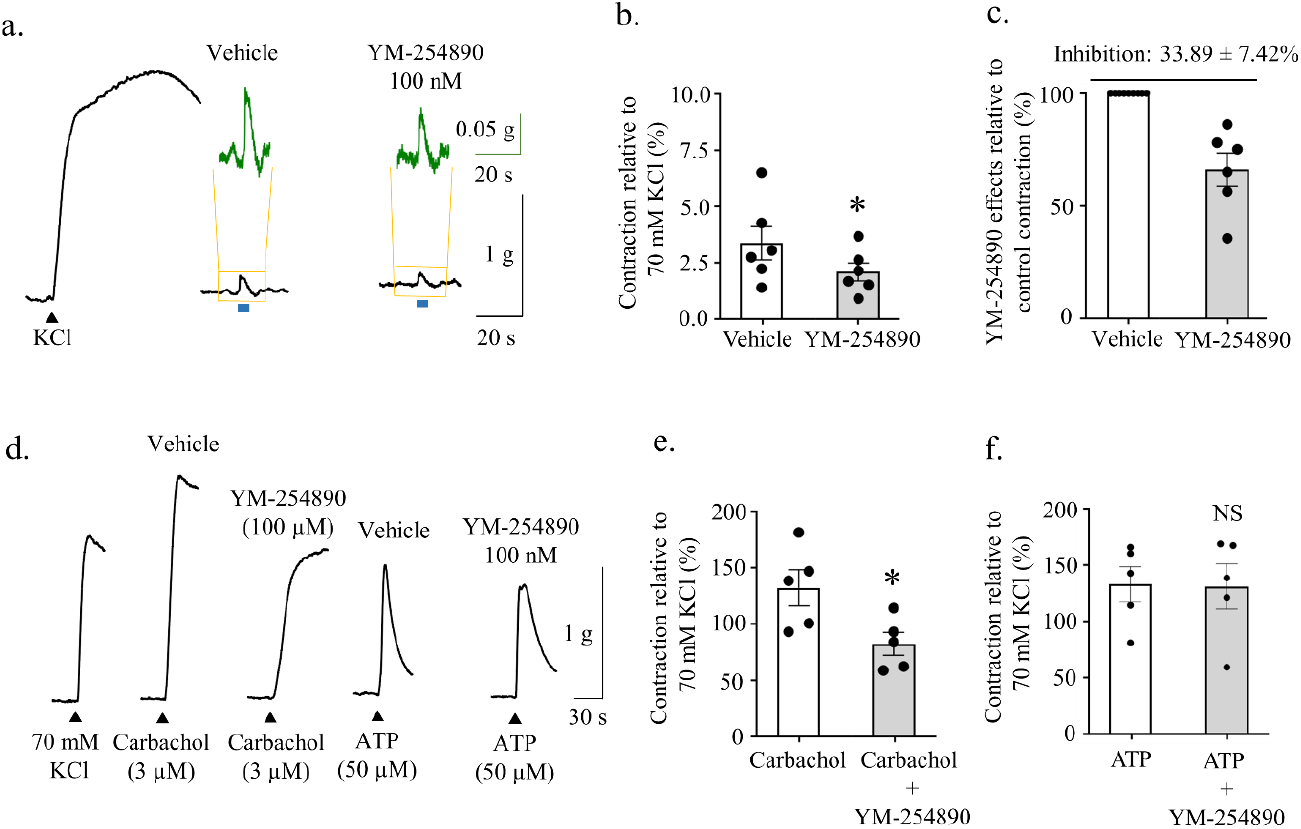
YM-254890 attenuates the urothelial cell-mediated local bladder contractions. (a) Representative recordings of the contractions evoked by 70 mM KCl and blue light (20 mW, 5 s). Blue light evoked urothelial cell-mediated local bladder contractions were recorded in the presence of vehicle and YM-254890 (100 nM). (b) Summarized data showing the inhibitory effects of YM-254890 (100 nM) on the urothelial cell-mediated local bladder contractions. (c) Summarized data of the percentage of inhibition of the urothelial cell-mediated local bladder contractions by YM-254890 (100 nM). (d) Representative recordings of the contractions evoked by 70 mM KCl and carbachol (3 µM) and ATP (50 µM). (e, f) Summarized data showing the inhibitory effects of YM-254890 (100 nM) on the carbachol (3 µM) and ATP (50 µM)-evoked contractile responses. The amplitude of the urothelial cell-mediated local bladder contractions was expressed as a percentage of the 70 mM KCl-evoked contractions in the same preparation. Each point represents the mean ± SEM (n = 6 for YM-254890 on the urothelial cell mediated contractions and n = 5 for YM-254890 on carbachol and ATP-evoked contractions). *Significantly different (p = 0.045) from the corresponding control values. Blue bars correspond to light stimulation. NS = non-significant (p = 0.860). Blue bars correspond to light stimulation.

It has been reported that acetylcholine-evoked muscarinic contractions in the detrusor smooth muscle require signaling through Gq_/11_ proteins^10^. On the other hand, it is not yet clear whether ATP-evoked purinergic contractions also need GPCR signaling to induce mouse bladder contractions. Therefore, we further investigated to specify the inhibitory effects of YM-254890 (100 nM) on the urothelial cell-mediated local bladder contractions, whether it is linked to muscarinic or purinergic signaling, using carbachol and ATP, respectively. YM-254890 (100 nM) significantly inhibited carbachol (3 µM)-evoked contractions but not ATP (50 µM)-evoked contractions (**Figure 3d, 3e, 3f**). The mean peak amplitudes of the carbachol (3 µM)-evoked contractions relative to 70 mM KCl-evoked contractions before and after the application of YM-254890 (100 nM) were 132.03 ± 16.11% and 82.39 ± 10.19% (n = 5) (**Figure 3e**), respectively. Whereas the mean peak amplitudes of the ATP (50 µM)-evoked contractions relative to 70 mM KCl-evoked contractions before and after the application of YM-254890 (100 nM) were 66.35 ± 7.89% and 65.50 ± 10.03% (n = 5) (**Figure 3f**), respectively. Our data indicates that GPCR signaling is essential for acetylcholine but not for ATP-evoked purinergic signaling pathways in mouse bladder. Taken together, these results suggest that inhibition of the urothelial cell-mediated local bladder contractions by YM-254890 (100 nM) could be due to the inhibition of acetylcholine-evoked GPCR signaling or somewhere else in this pathway that results in acetylcholine related contractions.

### Effects of Voltage-Gated Calcium Channel Inhibitor Nifedipine on the Urothelial Cell-Mediated Local Bladder Contractions

Nifedipine has been shown to inhibit L-type Ca^2+^ channels, thereby reducing bladder strip contractions by interfering with extracellular Ca^2+^ influx into the cells^15^. Signaling events of smooth muscle contractions are involved with Ca^2+^ influx through the voltage-gated Ca^2+^ channels and partly by intracellularly released Ca^2+16,17^. In this study, we sought to see whether voltage gated calcium channels are involved in the urothelial cell-mediated local bladder contractions. Nifedipine (2 µM) completely abolished the urothelial cell-mediated local bladder contractions (**Figure 4a, 4b, 4c**). These results suggest that the urothelial cell-mediated local bladder contractions require signaling pathways which involve Ca^2+^ influx through voltage-dependent Ca^2+^ channels.

**Figure 4.**
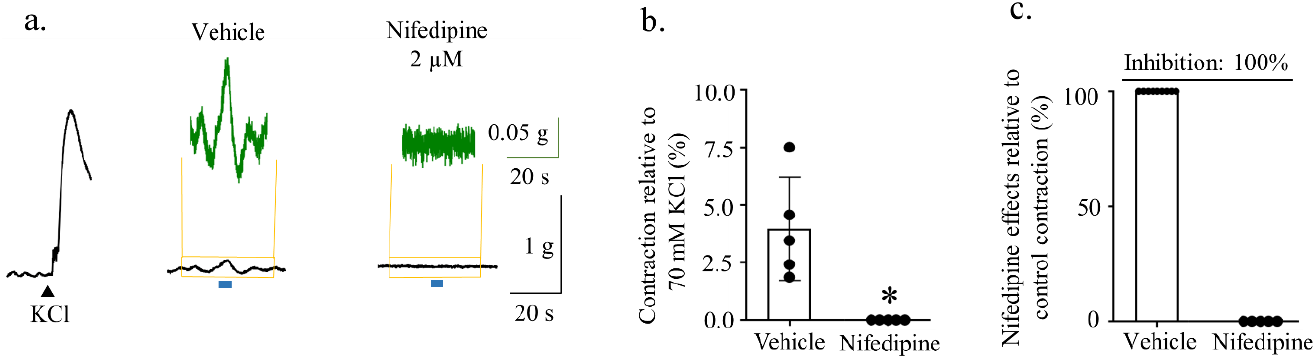
Effects of nifedipine, an L-type Ca^2+^ inhibitor, on the urothelial cell-mediated local bladder contractions. (a) Representative recordings of the contractions evoked by 70 mM KCl and blue light (20 mW, 5 s). Blue light evoked urothelial cell-mediated local bladder contractions were recorded in the presence of vehicle and nifedipine (2 µM). (b) Summarized data showing the inhibitory effects of nifedipine (2 µM) on the urothelial cell-mediated local bladder contractions. (c) Summarized data of the percentage of inhibition of the urothelial cell-mediated local bladder contractions by nifedipine (2 µM). The amplitude of the urothelial cell-mediated local bladder contractions was expressed as a percentage of the 70 mM KCl-evoked contractions in the same preparation. Each point represents the mean ± SEM (n = 5). *Significantly different (p = 0.016) from the corresponding control values. Blue bars correspond to light stimulation.

### Effects of Hemichannels Inhibitor Carbenoxolone Disodium and Exocytotic Molecules Release Inhibitor Brefeldin A on the Urothelial Cell-mediated Local Bladder Contractions

ATP can be released from urothelial cells by several mechanisms. The two main pathways are diffusion through hemichannels, and exocytosis^18,19^. We investigated whether these two important pathways are involved in releasing ATP or acetylcholine from the stimulation of the urothelial cells to produce the urothelial cell-mediated local bladder contractions. Here we used carbenoxolone disodium (CBX) and brefeldin A (BFA), which can block hemichannels and exocytosis pathways, respectively^20,21^.

CBX (100 µM) significantly inhibited the urothelial cell-mediated local bladder contractions (**Figure 5a, 5b**). The mean peak contractions relative to 70 mM KCl-evoked contractions were 2.18 ± 0.28%, which was reduced to 1.54 ± 0.20% (n = 6) by the addition of CBX (100 µM) into the bath solutions (**Figure 5b**). BFA (200 µM) also significantly inhibited the urothelial cell-mediated local bladder contractions (**Figure 5d, 5e**). The peak amplitudes of contractions relative to 70 mM KCl-evoked contractions were 2.29 ± 0.36%, which was reduced to 1.44 ± 0.26% (n = 5) after the addition of BFA (200 µM) into the bath solutions (**Figure 5e**). The percentage of inhibition by CBX (100 µM) and BFA (200 µM) were 33.89 ± 7.42% (n = 6) and 36.75 ± 6.20% (n = 5), respectively (**Figure 5c and 5f**). These results suggest that both hemichannels and exocytosis pathways are involved in releasing transmitter molecules from the stimulation of the urothelial cells.

**Figure 5.**
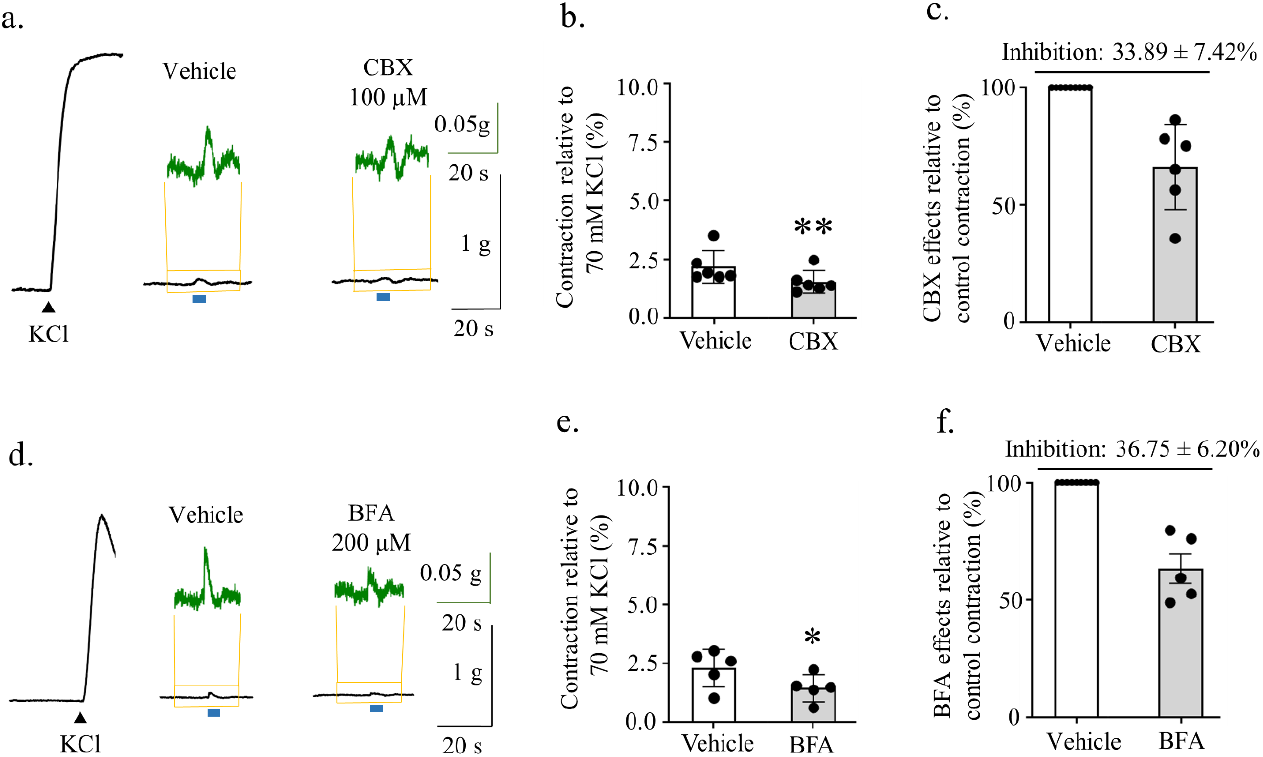
Effects of hemichannels inhibitor carbenoxolone (CBX) and exocytotic release inhibitor brefeldin A (BFA) on the urothelial cell-mediated local bladder contractions. (a and d) Representative recordings of the contractions evoked by 70 mM KCl and blue light (20 mW, 5 s). Blue light evoked urothelial cell-mediated local bladder contractions were recorded in the presence of vehicles and CBX (100 µM) and BFA (200 µM), respectively. (b and e) Summarized data showing the inhibitory effects of CBX (100 µM) and BFA (200 µM) on the urothelial cell-mediated local bladder contractions, respectively. (c and f) Summarized data of the percentage of inhibition of the urothelial cell-mediated local bladder contractions by CBX (100 µM) and BFA (200 µM), respectively. The amplitude of the urothelial cell-mediated local bladder contractions was expressed as a percentage of the 70 mM KCl-evoked contractions in the same preparation. Each point represents the mean ± SEM (n = 6 for CBX; n = 5 for BFA). Significantly different (**p = 0.005, and *p = 0.020) from the corresponding control values. Blue bars correspond to light stimulation.

## Discussion

The urothelial cells respond to sensory signals due to chemical and mechanical stimuli and release a variety of cell signaling molecules that are important for normal bladder function. We propose that these responses signal sensory neurons to convey mechanosensory information to the central nervous system. In the present study, we found that the urothelial cells, when optogenetically stimulated, can signal to non-neuronal cells to elicit bladder contractile activity (**Summarized in Figure 6**).

**Figure 6.**
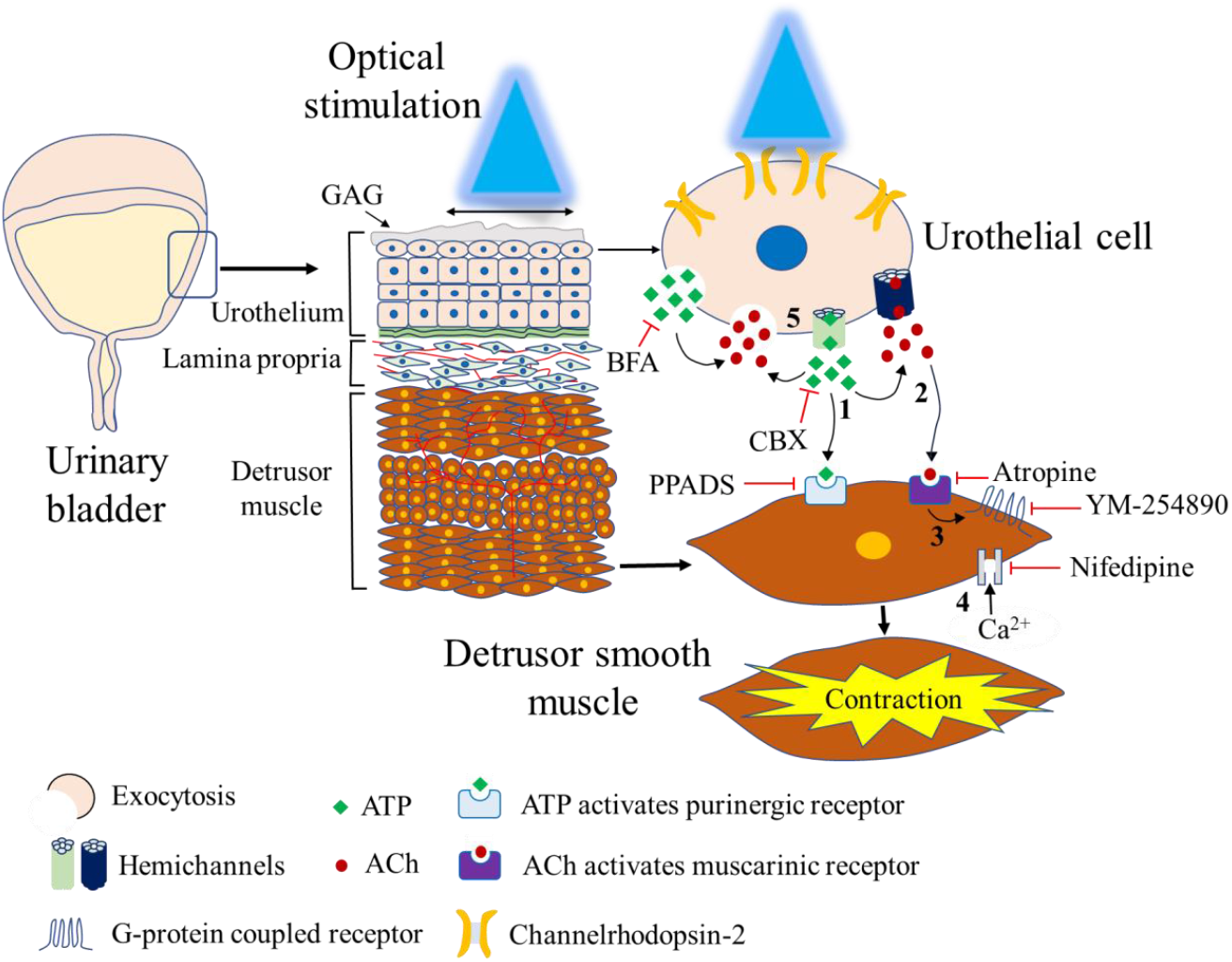
An outline of the predictive signaling events activated by stimulation of the urothelial cells leading to the urothelial cell-mediated local bladder contractions. **1)** The urothelial cell stimulation causes the release of ATP. Then the released ATP activates the purinergic signaling components on the detrusor smooth muscles. **2)** The released ATP also acts as a mediator of releasing ACh from the urothelium. The released ACh then activates the muscarinic signaling events on the detrusor smooth muscles. **3)** G-protein coupled receptors (GPCR) are involved in producing the urothelial cell-mediated local bladder contractions and it is related to the ACh activation of muscarinic receptors which thereafter activates the GPCR signaling to produce depolarization and contractions. **4)** The urothelial cell-mediated local bladder contractions signaling eventually causes the opening of voltage dependent Ca^2+^ channels to influx extracellular calcium into the cells. **5)** Stimulation of the urothelial cells releases ATP and ACh by means of exocytosis and hemichannels.

### Role of ATP in the Urothelial Cell-mediated Local Contractions

During the micturition cycle, stretching of the urothelium occurs, followed by the release of different signaling molecules such as ATP, ACh, nitric oxide, and substance P^2^. We first explored ATP and purinergic receptors because many different studies have suggested that ATP is the key molecule for the urothelial cell-mediated reflex bladder contractions^3,22,23^. Our data supports the findings of other studies demonstrating the importance of purinergic receptors in initiating the urothelial cell-mediated local bladder contractions (**Figure 2**).

We showed that PPADS at a concentration of 2 mM almost completely abolished the urothelial cell-mediated local bladder contractions (**Figure 2a-c**). The IC_50_ of PPADS for different P2X receptors varies from 1∼50 µM and P2Y receptors from ∼0.9 mM to 15 mM^24,25^. Therefore, while we don’t know the exact concentration at the receptor the 2 mM PPADS used in this study may have effects on P2Y receptors in addition to P2X. Additionally, the permeability of PPADS from the bladder bath solutions could influence its action. For instance, the application of atropine and ATP to the bath solution outside of the bladder did not inhibit the atropine-sensitive component of ATP-evoked contractions like as it did upon application within the bladder^5^. This finding suggests difficulty of the external absorption of these substances, which limits their effects from the external environment to the urothelium.

### Role of Acetylcholine (ACh) in the Urothelial Cell-Mediated Local Contractions

In *ex vivo* rat bladder preparation, it has been shown that muscarinic receptor inhibition by atropine significantly reduced the exogenous ATP-evoked contractions^5^. This result indicates that ATP may play a role in evoking cholinergic contractions of the bladder by influencing the release of ACh. In the present study, our data shows that muscarinic receptor inhibition with atropine significantly inhibited the urothelial cell-mediated local bladder contractions (**Figure 2**). This result was similar to the previous finding by Stenquist et al. 2017 which suggested that atropine-sensitive components of the urothelial cell-mediated local bladder contractions could be due to the influence of the predominant initial release of ATP from the urothelial cells leading to the release of ACh which activates the muscarinic receptors on the detrusor muscle^26^. Additional research is necessary to determine if the urothelial release of ATP and its atropine-sensitive role in the urothelial cell-mediated local bladder contractions can be altered in cases of bladder disease.

### Mechanisms of Releasing Transmitter Signaling Molecules from the Urothelial Cells

ATP is released from the urothelium by vesicular exocytosis and pannexin hemichannels in response to hydrostatic pressure or other stimulations^4,27^. In the present study, we provided direct evidence that the hemichannels inhibitor CBX significantly inhibited the urothelial cell-mediated local bladder contractions, indicating the involvement of hemichannels in releasing signaling molecules in response to local urothelial stimulation **(Figure 5a-c)**. Furthermore, exocytosis inhibitor BFA also caused significant inhibition of the urothelial cell-mediated local bladder contractions, suggesting exocytosis pathways are also involved **(Figure 5d-f)**. Unfortunately, with this data, we cannot specify the independent mechanisms for releasing ATP and ACh from the urothelium. Future studies are also needed to explore if there are any changes in the pathways that release transmitter molecules in response to urothelial stimulation in bladder diseases, as this has been hypothesized to be a critical component of diseases such as interstitial cystitis/ bladder pain syndrome^28^.

### Involvement of G-protein Receptor Signaling in Urothelial Cell-Mediated Local Contractions

We observed that urothelial cell-mediated local bladder contractions were significantly inhibited by a non-selective G-proteins inhibitor YM-254890 (**Figure 3**). Bladder contractions induced by muscarinic receptors are mediated via activation of Gq_/11_ proteins^10^. Jones et al., 2022 showed that YM-254890 (1 µM) abolished carbachol (200 nM) evoked bladder strips contractions^12^. However, in our study, YM-254890 (200 nM) was not abolished but significantly reduced the carbachol (3 µM)-induced contractions. This difference may be due to the relatively lower dose (1 µM vs 200 nM; 5 times lower) of YM-254890 we used to inhibit the higher dose (200 nM vs 3 µM; 5 times higher) of carbachol-evoked bladder contractions. Concentration-dependent inhibition of YM-254890 was also observed for the inhibition of GTPγS binding to Gq^29^. Furthermore, we found no significant inhibition of ATP-evoked bladder contractions by YM-254890. It has been proposed that ATP receptors in the bladder belong to two families: a P2X ion channel family and a P2Y G protein-coupled receptor family^30^. Bladder contractions are mediated predominantly via P2X receptors, while P2Y receptors mediate relaxation, which may be the possible reason for not inhibiting the ATP-evoked bladder contractions by YM-254890^31^.

### Calcium Signaling Events in Producing the Urothelial Cell-Mediated Local Contractions

An increase in intracellular cytoplastic Ca^2+^ concentration is fundamental to activate the contractile machinery either by Ca^2+^ influx or release of stored calcium from the intracellular sarcoplasmic reticulum^32,33,34^. In our present study, the urothelial cell-mediated local bladder contractions were abolished with calcium influx inhibitor nifedipine (**Figure 6**). Therefore, our data suggests that the urothelial cell-mediated local bladder contractions primarily depend on signaling pathways, which lead to an increase in intracellular Ca^2+^ levels through Ca^2+^ influx via voltage-dependent Ca^2+^ channels from extracellular stores.

### Study Limitations

In our study, we were unable to induce light-evoked distinguishable urothelial cell-mediated local bladder contractions from the mouse bladder strips (data not shown) and we were only able to evoke local contractions in the whole bladder preparations. This might be due to the decreased number of the urothelial cells in bladder strips, and their collective activation was not sufficient to produce a contraction. We also need to consider any potential damage to the urothelial cell layers and signaling potential when the bladder is cut into strips. With the whole bladder preparation, there is a sufficient population of cells available to activate the local contractions signaling pathway.

We showed that the urothelial cell-mediated local bladder contractions occur due to the release of ATP and ACh by activating ChR2 expressed into the urothelium **(Figure 2)**. We know that ChR2 activation can depolarize the cell and release ATP, similar to the activation of endogenous channels^3^. However, ChR2 is not normally expressed in the urothelial cells, and thus, the downstream signaling initiated by its activation may differ from endogenous cation channels like PIEZO1, TRPV4, or P2X2. This point needs to be considered when interpreting these results. Future studies could use newly engineered opsins that can mimic endogenous receptor activity and thereby link opsin activity to more specific downstream intracellular signaling cascades^35^.

## Conclusion

In conclusion, we have demonstrated that optogenetic stimulation of the urothelial cells can produce the urothelial cell-mediated local bladder contractions. This validates the previous hypothesis that the urothelial release factors can influence bladder contractions locally, without the need for signaling from the central nervous system. Further studies are needed to determine the relevance of these local signaling pathways in normal bladder physiology and pathophysiologic conditions.

## Data Availability

The data supporting this study’s findings are available to the corresponding author upon reasonable request.

## Acknowledgements

This study was supported by the Rita Allen Foundation Scholars Program Fund, Community Foundation of New Jersey and R01DK140233 (to A.D.M.); and 2022 Urology Care Foundation Research Scholar Award Program and Indian American Urological Association Sakti Das, MD Awards (to F.A.).

## Competing Interests

The authors declare no conflicts of interest, financial or otherwise.

## Author Contributions

F.A and A.D.M contributed to the conception and design of the study. F.A and K.J.R contributed to collecting and analyzing the data. F.A designed the figures and wrote the draft of the manuscript. A.D.M supervised the experiments, collected funds, interpreted the results, edited and reviewed the manuscript. F.A., K.J.R and A.D.M approved the final version of the manuscript.

## Notes

### Competing Interest Statement

The authors have declared no competing interest.

